# Reframing fundamental concepts of insect societies for bio-ontologies

**DOI:** 10.1101/2021.05.18.444717

**Authors:** Thiago S.R. Silva

## Abstract

Terminological and conceptual ambiguity surrounding core entities such as ‘colonies’ and ‘castes’ continues to hinder comparative analysis, data integration, and ontological interoperability in social insect research. While extensive work has addressed the descriptive and functional aspects of social organization, less attention has been paid to the ontological status of these entities and the consequences this has for formal representation in bio-ontologies. In this study, I provide a realist ontological analysis of ‘colonies’ and ‘castes’ grounded in distinctions between natural units, natural kinds, and realizable entities. I argue that insect colonies constitute bona fide biological individuals (natural units) demarcated through relations of causal unity and historical continuity, and therefore suitable for treatment as unified causal systems within ontology frameworks. By contrast, I show that ‘castes’ fail to satisfy the criteria for either natural units or natural kinds, due to their lack of independent causal cohesion, developmental plasticity, temporal variability, and context sensitivity. Treating ‘castes’ as autonomous entities or essentialized kinds introduces systematic instability and misrepresentation in bio-ontological models. To address these issues, I propose modeling ‘castes’ as realizable entities, specifically as roles instantiated in individual organisms and realized in biological processes under colony-regulated developmental and ecological conditions. This approach accommodates morphological canalization, phenotypic plasticity, temporal polyethism, and cross-taxon variability within a unified framework, while remaining consistent with the Basic Formal Ontology (BFO). My analysis clarifies longstanding conceptual ambiguities in the representation of social insect organization and provides a flexible, biologically faithful foundation for ontology design, supporting robust data interoperability and future empirical refinement.

## 1 Introduction

Terminological standardization has been a matter of debate among research groups that study insect societies for several years (Costa & Fitzgerald 1996, 2005; Peeters 2012; Neco et al. 2018; Silva & Feitosa 2019a; Sumner et al. 2018). According to several authors (e.g. Costa & Fitzgerald 2005; Crespi & Yanega 1995; Dew et al. 2016; Sumner et al. 2018), there are distinct reasons for the need for terminological univocity in this field of inquiry: (i) to provide a clearer conceptual landscape that enables researchers to explore drivers in the evolution of sociality in animals, (ii) to provide the linguistic components for more straightforward communication, whether textually or orally, and (iii) to provide a framework for unified concept representation and organization, which precedes implementations of formal specifications aimed at (meta)data interoperability, ultimately enhancing the Findability, Accessibility, Interoperability, and Reusability of (meta)data (see FAIR Guiding Principles, Wilkinson et al. 2016).

Aiming for concept clarity is an important first step for the standardization of specialized terminologies, although it does not directly entail standardization per se, since other terminological dimensions must be explored for consolidating shared sets of terms (Epstein 2012; Faber 2015; Faber & Léon-Araúz 2016). While the above-mentioned authors (and many others) provided distinct accounts for defining criteria for establishing unambiguous concepts of social categories in insect societies, few have explored how these categories can be translated into data models. While urging for ‘holistic’ approaches for defining and describing social categories, researchers generally opt for combined frames of reference when defining them, namely spatiostructural and functional frames (Michener 1974; Wilson 1975; Dew et al. 2016; Silva & Feitosa 2019a).

Although employing different frames of reference during concept representation can be considered the best approach for demarcating biological entities, researchers can equivocally define concepts through relations that violate one or more classification criteria in a given conceptual scheme, leading to inconsistent classifications. These inconsistencies can happen when researchers try to classify a set of entities using a *more-than-one* set of independent and not disjoint properties. Examples of inconsistent classifications in ‘caste’ representation include gamergates and ergatoids, as commonly understood in myrmecology (Peeters 1991; Peeters & Crozier 1988; Peeters 2019) (Figure 1). Traditionally, ‘castes’ in ants have been defined by a combination of properties belonging to two distinct frames of reference (i.e., spatiostructural and functional); workers are sterile apterous females (Figure 1.A) and queens are fertile macropterous females (Figure 1.B) (Höolldobler & Wilson 1990; Buschinger & Winter 1976, 1978; Peeters & Crozier 1988; Peeters 2019). In this particular example, gamergates would be subsumed in workers while ergatoids are subsumed in queens; additional defining properties, however, are conflicting with the properties of their supervening categories (Figure 1.C and D).

**Fig. 1.**
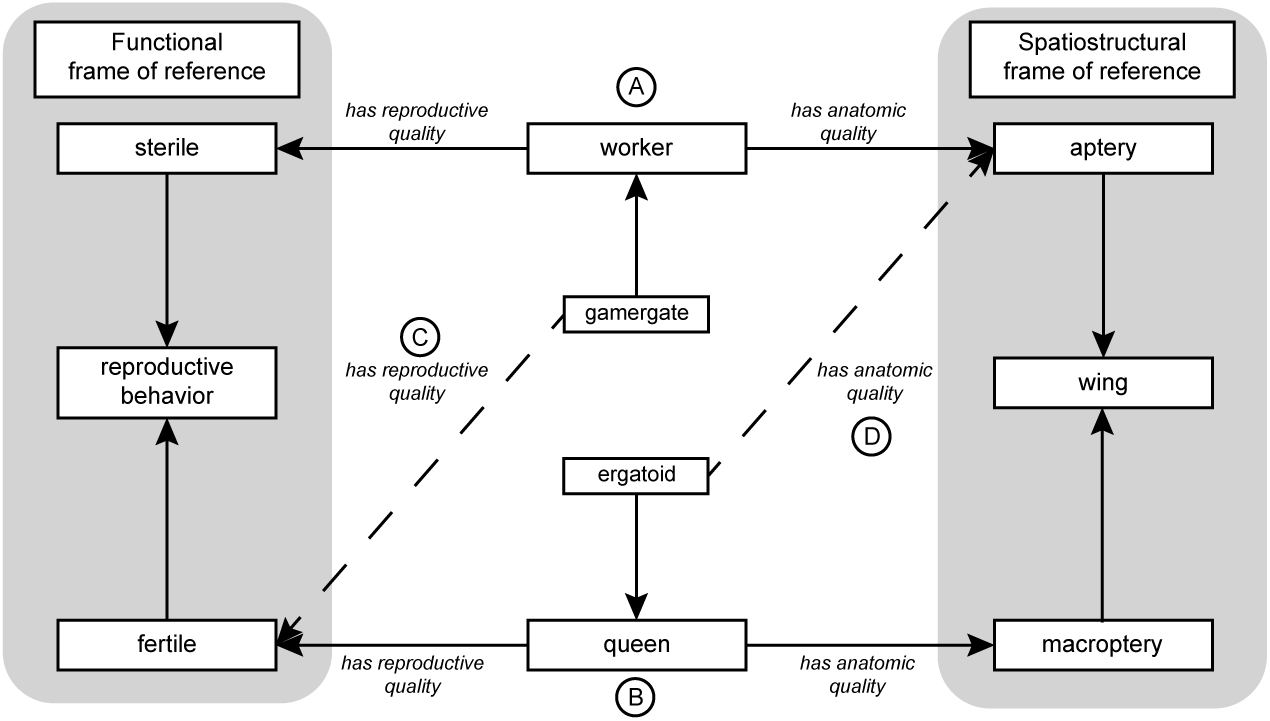
Example of ‘inconsistent classification’ in two ‘caste’ concepts applied in myrmecology: gamergate and ergatoid. Violations of classification criteria are marked as dotted lines. In this example, most relational properties are not necessarily compliant with formal ontologies (e.g., *incapable of*, *determined by presence of*, *has anatomic quality* ) and are used only for didactic purposes.

Although the inconsistencies discussed here appear terminological or empirical, they point to a deeper ontological problem concerning how biological entities are individuated and classified. In biological ontology design, a crucial distinction must be drawn between partitioning and classification, since they involve different kinds of epistemic activities and result in different types of entities. Partitioning aims to demarcate natural units — individual entities that can be identified through their internal connectedness — whereas classification presupposes partitioning and aims to identify natural kinds, or classes of entities, based on shared defining properties. Conflating these two epistemic activities leads to systematic errors in ontology design, particularly when collections of organisms are treated as individuals or when context-dependent properties are reified as kinds. Recognizing this distinction is essential for the consistent representation of biological concepts in ontology frameworks.

This work aims to provide a logical account for translating two fundamental concepts in the study of social insects to a realist ontology — i.e., ‘colony’ and ‘caste’ — which can be used by researchers for describing and organizing entities belonging to non-human societies, as well as for the provision of evidential criteria for evaluating constitutive explanations of ‘social’ entities. In doing so, I will argue that these two concepts correspond to different ontological categories: individual ‘colonies’ can be treated as natural units identified through relations of causal unity (i.e., through partitioning), whereas ‘castes’ are better understood as realizable entities — specifically as roles borne by individual organisms within a ‘colony’ context. This interpretation reflects the fact that caste-related properties (e.g., reproductive activity, foraging, brood care) are only manifested under certain biological and temporal conditions and are thus realized in processes rather than being enduring kinds.

I will argue that both concepts can be demarcated under two common frames of reference used in biological sciences, according to Vogt (2019a): functional and historical/evolutionary. I will adjust and refine previous propositions for a ‘caste’ ontology while addressing some misconceptions made by Silva & Feitosa (2019a). By reinterpreting ‘castes’ as realizable entities, this work seeks to provide a more flexible and ontologically consistent foundation for representing insect social categories in bio-ontologies, accommodating both functional and temporal variation in ‘caste’ expression. Lastly, I will provide a brief account of the limitations of current approaches to demarcate subcategories of ‘castes’.

## 2 Methods

### 2.1 Specialized terms

In this work, I used quotation marks when referring to the names ‘colony’ and ‘caste’, in order to highlight the inappropriateness of their usage for communication in contemporary biology [cf. discussion in Herbers (2007, 2020) and Sleigh (2004)]. I understand that reevaluating the linguistic component of terminological units — i.e., multidimensional units composed of linguistic, cognitive, and situational components (Cabre 2003) — is essential for the development of a more inclusive and diverse science. I suggest that a reevaluation of this magnitude should be conducted by a diverse group of researchers through a transdisciplinary approach, which is a desirable condition for addressing linguistic modifications. Since it is not the aim of the present work to reevaluate the historical usage of these names within the discipline, I refrained from providing alternative terms to the concepts that refer to aggregations of individual organisms belonging to groups of social insects and to distinct categories of phenotypes in non-human societies, lest I replace an oppressive-laden term with another.

### 2.2 Frames of reference and relations of causal unity

The present analysis is conducted within a realist ontological framework as operationalized in the Basic Formal Ontology (BFO). Accordingly, the critiques and modeling choices developed here are relative to specific commitments concerning mind-independent biological individuals, causal unity, and the use of formal ontology as a tool for representing biological reality. This methodological stance does not presuppose that such commitments exhaust all legitimate perspectives on biological organization; rather, it reflects the fact that BFO-based realism currently provides the dominant upper-level framework for ontology development in biodiversity and phenomics, and therefore constitutes a relevant and widely adopted target for critical analysis.

Within this framework, I employed realist-based approaches to explore how fundamental concepts from the study of social insects can be adapted for application in bio-ontologies. The primary goal was to develop extensions of existing ontological entities that remain conformant with BFO while addressing well-known challenges in the identification of biological individuals. In particular, I focused on two interrelated problems: (1) the granularity dependence of bona fide boundaries and (2) the perspective dependence of bona fide boundaries. Both issues concern discontinuity (and, conversely, connectedness) as well as qualitative heterogeneity across biological scales (Vogt 2019b).

To address these challenges, I modeled ‘colonies’ as natural units demarcated through specific types of causal unity, each corresponding to distinct frames of reference (Smith et al. 2015; Vogt 2019a). Two frames of reference were employed: functional and historical/evolutionary. Within these frames, the internal connectedness underpinning a colony’s status as an individual biological unit is grounded in patterns of functional integration among organisms and in relations of common descent and historical continuity. Under this interpretation, ‘colonies’ can be treated as aggregates of functionally and historically integrated organisms that maintain identity through continuous causal relations among their members. Such identity may be transiently disrupted during processes such as fission or budding without immediately negating unit status, provided that causal and historical continuity is preserved.

To further clarify the treatment of non-structural boundaries as mind-independent within a realist framework, I followed the account of functional and historical/evolutionary units proposed by Vogt et al. (2012). Functional units can be identified through (i) dispositions independent of morphogenesis and (ii) dispositions involving morphogenetic processes, whereas historical/evolutionary units are identified through (iii) common causal history, referring to structural integrity and persistence over time, and (iv) common historical origin, referring to constituent historical relations among entities distributed across time and space. These criteria provide a principled basis for demarcating colonies as biological individuals within a BFO-aligned ontological framework, while remaining explicit about the frames of reference under which such demarcations are made.

### 2.3 Conceptual dependence and realizability

In considering ‘castes’, I adopted BFO’s framework of *realizable entities*. Realizable entities are specifically dependent continuants that inhere in independent continuants and are realized in processes (Arp et al. 2015). Unlike material entities, they do not exist continuously but are manifested only under specific biological conditions. Within this framework, I evaluated whether ‘caste’ entities can be consistently modeled as realizable entities — particularly as *roles* — that depend on the social and developmental context of the organism.

Roles, in BFO, are realizable entities that exist because their bearers are embedded within specific intrinsic and relational contexts and are capable of realizing certain functions under appropriate conditions. In this context, bearing a ‘caste’ role should not be understood as merely possessing an abstract developmental potential. Rather, role bearing is grounded in biologically realized capacities that arise from concrete developmental, physiological, or morphological organization. For instance, morphological differentiation associated with reproductive or worker pathways provides empirical evidence that an organism bears a corresponding ‘caste’ role, even if that role is not currently realized in an ongoing process. Role realization, by contrast, requires the occurrence of specific biological activities (e.g., oviposition, brood care, foraging) under appropriate social and environmental conditions. This distinction preserves empirical tractability while preventing the conflation of dispositional capacity with taxonomic identity.

Modeling ‘castes’ as roles rather than as natural kinds also accommodates the developmental and contextual flexibility observed in many social insects, while remaining consistent with BFO’s treatment of realizable entities. Because ‘caste’ differentiation often occurs during post-embryonic development, individuals may bear multiple unrealized potentialities that become manifest under specific environmental or epigenetic conditions. For example, an ant larva may develop into a ‘queen’ or a ‘worker’ depending on the realization of underlying developmental dispositions regulated by social and physiological factors. This interpretation captures the dynamic and conditional realization of ‘caste’ roles across the life cycle of social insects, integrating ontogenetic, social, and functional dimensions of ‘caste’ expression. It thereby avoids the categorical rigidity of kind-based classifications, aligns with ontological realism as prescribed in the BFO, and provides a flexible foundation for integrating ‘caste’-related data into bio-ontologies in a manner consistent with existing representations of functions, dispositions, and processes.

Building on this conceptual framework, I apply the realizable entity interpretation of ‘castes’ to representative insect societies, illustrating how these roles can be formally modeled and operationalized within bio-ontologies.

### 2.4 OWL formalisms

To improve readability and accessibility for readers not familiar with ontology engineering or the Web Ontology Language (OWL), this section summarizes the formal notation used throughout the paper. The axioms presented in the Results and Discussion follow standard Description Logic–style OWL syntax as employed in BFO-aligned bio-ontologies. Table 1 summarizes the small set of OWL-style symbols and BFO relations used in the formal axioms presented in this paper.

**Table 1.** Summary of the OWL-style formal notation employed for ontological representation in this study.

| Notation | Explanation |
| --- | --- |
| $A \sqsubseteq B$ | Subclass relation: every instance of class $A$ is also an instance of class $B$. |
| $A \equiv B$ | Class equivalence: classes $A$ and $B$ have exactly the same instances. |
| $\wedge$ | Logical conjunction ("and"): all stated conditions must hold simultaneously. |
| $\vee$ | Logical disjunction ("or"): at least one of the stated conditions must hold. |
| *inheres_in* | BFO object property relating a specifically dependent continuant (e.g., a role) to its bearer (e.g., an organism). |
| *realized_in* | BFO object property relating a realizable entity to the process in which it is realized. |
| **role** | A BFO realizable entity that is context-dependent and not essential to the kind of its bearer. |

These notational conventions are used to express minimal formal commitments that serve as conceptual witnesses of the proposed ontological distinctions throughout the text, rather than as exhaustive or prescriptive data models.

## 3 Results

### 3.1 Natural kinds

In philosophical and scientific discourse, a “natural kind” is understood as a grouping of entities that reflects real divisions in nature rather than human-imposed classifications. The idea is that such groupings correspond to objective, mind-independent structures in the world and not merely to convenience, human interests, or historical contingencies.

Classic examples often invoked as paradigms of natural kinds come from chemistry (such as the chemical elements) but natural-kind classifications also appear in physics (fundamental particles), astronomy (types of galaxies), and, more controversially, biology. The claim behind natural-kind realism is that our best scientific classifications aim to discover and reflect these natural divisions, rather than impose conventional categories guided by human motivations.

Philosophers have proposed several criteria commonly associated with natural kind membership. First, members of a natural kind should share certain (typically natural, often intrinsic) properties, though sharing a property alone is not sufficient for kind-hood. Second, natural kinds should support inductive inferences: knowing that one individual belongs to a kind should permit predictions or generalizations about others. Third, many accounts suggest that kinds, to qualify as natural, should participate (or at least consistently correlate) with stable laws or regularities.

However, the notion of natural kinds is not uncontroversial. In fields with high biological, developmental, or evolutionary variability (such as biology) it becomes difficult to identify kinds fulfilling all criteria consistently. For example, what counts as a “species”, “subspecies”, or morphological variety may shift over time, and traits may vary under environmental or developmental influences. Such variability challenges essentialist views that a natural kind must have stable, necessary, and sufficient defining properties. As a result, many philosophers have proposed more flexible accounts (such as “cluster-kind realism”) where a natural kind is defined not by a single essential property but by a cluster of properties that tend to co-occur because of underlying causal or homeostatic mechanisms.

Given these philosophical debates, whether a given biological grouping (e.g., a ‘caste’ of social insects) qualifies as a natural kind depends critically on whether it meets these criteria: sharing stable, relevant properties; supporting reliable inductive inferences; and exhibiting sufficient stability across contexts. Importantly, failure to qualify as a natural kind does not entail failure to qualify as a natural unit, and vice versa; the two notions impose distinct ontological requirements. As I will show in subsequent sections, the variability, plasticity, and context-sensitivity common in ‘caste’ phenomena complicate the attribution of natural-kind status to ‘castes’.

### 3.2 Natural units

The distinction between bona fide and fiat boundaries is essential for identifying natural units (bona fide objects) versus artifactual entities (fiat material entities). Bona fide objects possess continuous, intrinsic boundaries, whereas fiat material entities possess imposed or arbitrary boundaries (Smith and Varzi 1997a, 2000; Smith 1994, 1995, 2001). Recognizing this distinction is crucial not only for identifying natural units but also for organizing, structuring, and managing domain knowledge. Top-level categories of bona fide objects have been used to organize the material realm according to compositional hierarchies (e.g., Woodger 1929; Novikoff 1945; Oppenheim and Putnam 1958; Simon 1962; Wimsatt 1976, 1994; Mayr 1982; Eldredge 1985; Riedl 1985; Levinton 1988; Craver & Bechtel 2007; Winther 2011; Vogt 2017, 2019a).

In the identification of natural units, two conceptual challenges arise. First, *discontinuity* (or, conversely, connectedness) emerges from the granularity dependence of bona fide boundaries: at different scales of observation, continuous entities may appear fragmented or merged. Second, *qualitative heterogeneity* arises from perspective dependence: the same entity may exhibit different properties when analyzed under distinct frames of reference (e.g., spatiostructural, functional, or historical/evolutionary). Both challenges complicate the consistent demarcation of natural units.

The notion of *causal unity* offers a solution to these challenges. By focusing on the causal relations that maintain an entity as a cohesive whole, one can identify natural units despite local discontinuities or heterogeneous properties. Causal unity allows the integration of multiple frames of reference, ensuring that a bona fide object can be recognized as a single, coherent entity even when its components display different structural, functional, or historical characteristics (Vogt 2019a). In this way, causal unity provides a principled criterion for partitioning entities as natural units, which can then be linked to functional and historical/evolutionary frames of reference in subsequent analyses.

### 3.3 ’Colonies’ of social insects as unified causal systems

Insect ‘colonies’ are aggregates of conspecific organisms that occupy a given geographical space and share kinship, meaning that members typically exhibit high genealogical relatedness and live in close proximity (Strassmann & Queller 2010). Each individual belonging to a ‘colony’ is considered a member part of this aggregate, and, thus, its membership relation establishes the ‘colony’ as an aggregation of organisms. Members are causally connected to one another through relations of *causal unity*, which define the colony as a natural unit independent of human partitioning activities.

Some studies have approached ‘colonies’ under the complex adaptive systems (CAS) paradigm (Bonabeau 1998; Bonabeau et al. 1998), where properties at coarser levels emerge from interactions among components at finer levels and may feedback to influence subsequent interactions (Levin 1998). While useful for explaining certain spatiotemporal phenomena — such as trails, nests, or bivouacs emerging from non-morphogenetic dispositions (Bonabeau et al. 1998; Mlot et al. 2011; Phonekeo et al. 2017; Franks 1989) — CAS approaches are limited in accounting for the historical/evolutionary properties of colonies.

To address these limitations, I model ‘colonies’ as *causally relatively isolated entities*— i.e., unified causal systems structured through, and maximal relative to, specific types of causal unities (Ingarden 1983; Smith & Brogaard 2003). Two types of causal unity contribute to the connectedness of a colony: (i) causal unity via bearing a specific function; and (ii) causal unity via common historical/evolutionary origin (Vogt 2019a).

Colonial connectedness is maintained through properties that promote internal homeostasis (Schmickl & Karsai 2018). Key properties include chemical space, self-organization, and resilience. These properties can be linked to specific frames of reference, and their constituent parts can be further partitioned within their respective perspectives. The following subsections elaborate on how these properties relate to the functional and historical/evolutionary frames of reference of ‘colonies’.

#### 3.3.1 Functional frame of reference and functional properties of ‘colonies’

Historically, structural properties of ‘colonies’ have often been associated with the physical structures they occupy, such as nests (Jones et al. 2004). However, the structural properties of ‘colonies’ differ substantially from those of their physical niches, though the latter provide support for maintaining connectedness.

In the case of social insects, nests and “living structures” are not considered physical coverings in the strict sense of Smith et al. (2015), since they are not composed solely of an enclosing physical structure. Instead, they can be understood as *niche tokens* (Smith & Varzi 1999), often conceptualized as double-hole structures (Smith & Varzi 2001; see also Casati & Varzi 1994). These consist of: (i) a central hole representing the ‘colony’ proper (tenant) and its immediate medium (air or other material displaced to fit members), (ii) a surrounding medium (environing hole; e.g., chambers, tunnels, floors, or trails) for movement, respiration, and other activities, and (iii) often an external enclosing structure (retainer) surrounding both holes.

The surrounding medium is an important component that allows for connectedness of the ‘colony’ by supporting a variety of causally-linked processes. For example, it enables thermoregulation: in army ants, bivouac core temperatures are maintained above ambient levels through collective respiratory activity, promoting rapid development of immature stages and long-term colony stability (Franks 1989; Bartholomew et al. 1988). In this particular case, the crevices that compose the core of the ‘colony’ (i.e., the spaces between each individual organism) are filled by a mixed medium (i.e., air plus the metabolic products of respiratory processes) which provides the ideal thermal conditions for maintaining their respective tenants. The mixed medium also allows chemical interactions via volatile compounds such as pheromones, providing functional connections between colony members. Despite volatile compound components being structurally bound (normally in the form of weak interactions) to one another, the chains of production and recognition of these compounds functionally link individual organisms, resulting in colony-level coordinated behaviors such as acceptance, aggression, or exclusion.

Chemical signatures play a major role in functional connectedness. Cuticular hydrocarbons vary by ‘caste’ and sex, and specific patterns are recognized by colony members to guide behavior (Oi et al. 2015). They are a major force promoting functional connectedness of insect aggregations; specific patterns of chemical structure (i.e., chemical profiles) are recognized by the members of a given colony and these members decide how they must respond when faced with this variation (whether in an aggressive manner or not). If the pattern of the ‘colonial’ chemical structure is dissimilar enough, or if its concentration is lower than a given threshold, disconnection unfolds through aggressive behaviors or exclusion of the member from the ‘colony’. Even in cases where a ‘colony’ is considered ”fragmented”, - such in polydomous ‘colonies’, where members occupy spatially separated nests - shared chemical space maintains functional unity.

’Colonies’ are dynamic systems; functional connectedness may fluctuate as individuals temporarily leave the pheromone space or move within the environment. This limitation is accounted for by resilience mechanisms, which restore functional cohesion over time (see Results 3.3.2).

Self-organization is another key attribute supporting functional connectedness. Dispositional properties of individuals promote spatial and social structuring in response to local environmental or ecological constraints. Both spatial organization (in the form of movement and density) and social organization (in the form of task allocation) are fundamental for the regulation of a ‘colony’. Spatial organization is believed to be an important mechanism for colonial regulation since it allows for the aggregate to circumvent pitfalls of excessive growth during extremely favorable environmental conditions, or to avoid colonial collapse when faced with sub-optimal or catastrophic situations. In this sense, spatial organization is intrinsically related to social organization, since it is believed that interaction rates are typically density-dependent, meaning that individual movement, ‘colony’ size, and ‘colony’ density significantly alter group dynamics (Pacala et al. 1996).

An example of self-organization is trail formation in ants. When a food source is discovered, the first forager recruits others, increasing forager numbers until the colony reaches a stable population at the food site. Positive and negative feedback loops govern this process, and the system stabilizes when the recruitment rate balances the abandonment rate. This dynamic regulation of individuals reflects a colony-level disposition that is independent of morphogenesis.

Finally, movement zones or spatial fidelity zones also contribute to functional connectedness in insect societies (Sendova-Franks & Franks 1995; Modlmeier et al. 2019). They have been linked to an individual’s behavioral repertoire and are responsible for guiding them to specialized tasks, reducing task-switching costs, and for supporting ‘colony’ stability despite demographic or environmental fluctuations (Baracchi & Cini 2014; Heyman et al 2017; Jandt & Dornhaus 2009; Mersch et al 2013; Powell & Tschinkel 1999; Robson et al. 2000; Seeley 1982). Together, chemical communication, self-organization, and spatial fidelity allow ‘colonies’ to maintain functional unity as natural units over time.

#### 3.3.2 Historical/evolutionary frame of reference and historical/evolutionary properties of ‘colonies’

Under a historical/evolutionary frame of reference, the connectedness of ‘colonies’ is maintained through relations grounded in common descent and shared genealogical history. In this view, the integrity and persistence of a ‘colony’ emerge not only from its functional connectedness at a given time, but also from its continuity through successive generations of individuals that share common kinship. Thus, ‘colonies’ can be understood as historical individuals unified by causal relations of descent that connect their parts through time.

Self-recognition and kin-recognition are central mechanisms maintaining this historical connectedness. Members of a ‘colony’ recognize one another through a combination of visual and chemical cues that signal common ancestry or shared genetic makeup (Strassmann & Queller 2010). Recognition systems minimize intracolony conflict and promote cooperation, ensuring that individual behaviors align with ‘colonial’ demands. When recognition fails — such as in cases of usurpation or parasitism — the breakdown of genealogical coherence can result in the disintegration of the colony as a natural unit.

In social insects, recognition often relies on genetically encoded chemical cues present on the cuticle. These cues are shaped by both genetic relatedness and shared environmental influences, producing a characteristic colony odor (van Zweden & d’Ettorre 2010). For example, in honey bees, workers can distinguish nestmates from non-nestmates through slight variations in hydrocarbon profiles derived from common ancestry (Breed 2014). However, nestmate recognition based on colony odors is not restricted to vertically acquired pheromones, but can also be influenced by horizontally acquired pheromone profiles. In *Reticulitermes speratus*, for example, nestmate recognition based on gut symbiont profiles, which is primarily transmitted vertically from parents to offspring but secondarily influenced by horizontal acquisition from shared food sources (Matsuura 2001), can reflect both a functional unity via shared microbial processes and a historical continuity of microbial and host lineages. Such examples illustrate how kin recognition can function as (but is not restricted to) a causal mechanism linking present members of a colony to their ancestral lineage.

Colony demography, another property linked to historical connectedness, depends on the modulation of reproductive and developmental pathways that maintain genealogical integrity. Just as recognition mechanisms mediate continuity between present and ancestral members, demographic regulation ensures the long-term persistence of lineage-defining causal factors. Through the regulation of ‘caste’ ratios, replacement of reproductives, or production of dispersing alates, colonies maintain the continuity of their genetic lineages. For instance, in termites, when a queen dies, replacement reproductives (neotenics) may arise from within the same colony, preserving genealogical continuity (Vargo et al. 2012; Vargo 2019); in other cases, new reproductives found descendant colonies that inherit genetic and chemical features from the parent colony (Vargo 2019). In both scenarios, causal unity via common descent is maintained at the level of the ‘colony’ or broader “metacolony” lineage.

Common kinship also provides the basis for adaptive responses to ecological pressures. ‘Colonies’ with high relatedness tend to exhibit greater cohesion in cooperative tasks such as brood care, defense, and nest construction, because their genealogical structure aligns the reproductive interests of their members. Conversely, genetic heterogeneity — introduced, for example, through multiple mating or adoption of unrelated individuals — can generate internal conflicts over reproduction and resource allocation, potentially destabilizing the colony (Ratnieks et al. 2006). Thus, genealogical coherence acts as a stabilizing force that maintains the ‘colony’s’ integrity across its lifespan.

From an ontological perspective, ‘colonies’ structured through common kinship relations exemplify causal unity via common historical/evolutionary origin (Vogt 2019a). Their connectedness arises from a web of causal relations that trace back through reproduction and descent, linking present members to a shared origin. In this sense, the ‘colony’ can be regarded as a historical individual, i.e. a temporally extended system whose identity is preserved through causal continuity of its constituent lineages and their genealogical relations.

### 3.4 Limitations of treating ‘castes’ of social insects as natural units and natural kinds

Considering the accounts of ‘colonial’ connectedness presented in the previous sections, ‘castes’ can be understood not as independent units but as properties emerging from the developmental and regulatory architecture of a given colony. ‘Colonies’ possess developmental dispositions, which are rooted in their gene pool and modulated by ecological and social conditions, to produce individuals expressing certain phenotypes in response to intrinsic and extrinsic stimuli. On this view, ‘caste’ differentiation reflects the realization of these ‘colony’-level dispositions at the organismal level, conditioned by the ‘colony’s’ internal state and environmental context, rather than the manifestation of autonomous, colony-level units.

This perspective has important consequences for the ontological status of ‘castes’. Although ‘caste’ differentiation contributes to ‘colony’ organization by channeling organism-level dispositions (e.g., reproductive capacity, foraging behavior, defensive morphology), ‘castes’ themselves do not satisfy the criteria required of natural units. Unlike ‘colonies’, they lack causal, functional, spatial, and historical coherence as integrated biological individuals. Rather than constituting unified causal systems, ‘caste’ categories merely collect organisms that share certain properties at particular times and under particular conditions.

This becomes evident when considering how ‘caste’ categories are empirically identified in studies of social insects. Organisms within a ‘colony’ may share phenotypic or behavioral traits that motivate labels such as “worker”, “soldier”, or “queen”, yet these traits vary widely across ‘colonies’, species, and ecological contexts. Moreover, similar traits recur across organisms belonging to different ‘colonies’. This cross-‘colony’ recurrence encourages the treatment of ‘castes’ as natural groupings, but such groupings do not correspond to cohesive biological entities in the same sense as ‘colonies’ do.

Despite this, researchers often treat both ‘colonies’ and ‘castes’ as terms that directly refer to entities in biological reality. Under this assumption, the corresponding entities are taken to be natural kinds, their mental representations are concepts, and the associated linguistic expressions are kind terms (Vogt et al. 2022). Accordingly, ‘caste’ concepts are frequently defined intensionally using genus–differentia structures (e.g., “a queen is a reproductive female”). While such definitions provide semantic clarity, they rarely support reliable identification of instances in nature. Diagnostic criteria remain incomplete, and the application of ‘caste’ terms often proves inconsistent across taxa, developmental systems, and ecological conditions.

These conceptual difficulties are mirrored and amplified in ontological modeling practice. When ‘castes’ are treated as natural kinds, defining properties must be attributed to them. Yet the pronounced variability, developmental plasticity, and taxon-specificity of ‘caste’ expression undermine the assumption that such properties are stable or essential. As a result, treating ‘castes’ as natural kinds introduces systematic instability into bio-ontological representations.

The limitations of this approach become particularly clear when ‘castes’ are formalized within bio-ontological frameworks. When modeled as natural units analogous to ‘colonies’, ‘castes’ are implicitly treated as causally unified biological individuals composed of multiple organisms. A naive but tempting formalization would therefore assert axioms such as:

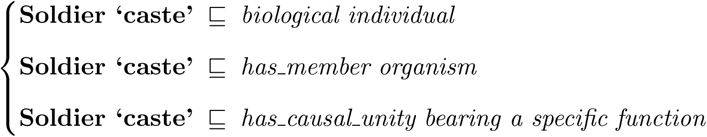

Such representations commit the ontology to the existence of higher-level biological individuals (i.e., ‘castes’) with their own integrity conditions and causal unity. However, as argued above, collections of organisms sharing a ‘caste’ label do not exhibit the spatial, temporal, functional, or historical connectedness required of natural units. Soldiers belonging to the same ‘caste’ may be spatially dispersed, temporally asynchronous, or developmentally heterogeneous, and they participate in unified causal processes only through their integration within the ‘colony’. Treating ‘castes’ as natural units therefore results in a misattribution of causal unity and the postulation of biological individuals lacking independent ontological grounding.

A second, more common strategy treats ‘castes’ as natural kinds defined by essential properties, often following a genus–differentia pattern. For example, a ”queen” may be modeled as a subtype of organism characterized by reproductive capacity:

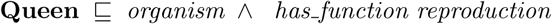

or, more strongly, as an equivalence class:

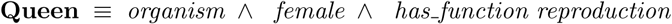

These axioms are presented schematically; in a fully BFO-compliant representation, reproductive capacity would itself be modeled as a realizable entity inhering in the organism and realized in reproductive processes.

While such definitions appear ontologically precise, they fail to accommodate the empirical variability of ‘caste’ expression across taxa and developmental contexts. Reproductive capacity may be transient, suppressed, or distributed across multiple phenotypes (e.g., gamergates, worker-laid eggs, ergatoid queens), and individuals may shift between functional states over their lifespan. Under essentialist modeling strategies, these dynamics force either repeated reclassification of organisms or the proliferation of increasingly ad hoc subclasses, undermining both taxonomic stability and semantic interoperability.

These problems are further exacerbated in comparative contexts. A single ”Worker” class defined by properties such as non-reproductivity or foraging behavior cannot be consistently applied across systems with fixed morphological ‘castes’ (e.g., *Pheidole*), temporal polyethism (e.g., *Apis mellifera*), or developmentally flexible worker phenotypes (e.g., termites). Consequently, the same class label comes to denote biologically heterogeneous entities grounded in incompatible developmental architectures, eroding the explanatory and integrative value of the ontology.

From an ontological standpoint, this contrast is fundamental. ‘Colonies’ correspond to independent biological units with cohesive causal and historical identity, making them plausible candidates for natural-kind-like treatment. ‘Castes’, by contrast, correspond more naturally to realizable entities in the sense of BFO, that is, dispositions, functions, or roles instantiated in organisms under specific developmental and environmental conditions. As such, ‘castes’ lack the autonomous identity, historical persistence, and causal unity characteristic of natural kinds.

Taken together, both the conceptual analysis and the formal modeling examples demonstrate that treating ‘castes’ as natural units or natural kinds leads to systematic ontological misalignment. In the former case, causal unity is incorrectly attributed to collections of organisms lacking independent cohesion; in the latter, essential properties are imposed on phenomena that are intrinsically context-dependent, temporally variable, and developmentally plastic. These considerations support the conclusion that ‘castes’ are better understood not as autonomous biological entities, but as context-dependent realizations of organism-level capacities within colony-regulated developmental regimes.

This shift in perspective motivates an alternative ontological treatment of ‘castes’ that focuses on the types of properties they instantiate, specifically, realizable entities governing functional, developmental, and ecological roles. In the following subsection, I show how modeling ‘castes’ as realizable entities accommodates context-sensitivity, developmental plasticity, and cross-taxon variability while avoiding the categorical problems associated with treating them as natural units or natural kinds.

### 3.5 ‘Castes’ of social insects as realizable entities

In light of the limitations of treating ‘castes’ as natural units or natural kinds, a more appropriate ontological interpretation situates ‘castes’ within the BFO category of realizable entities. This framework accommodates the developmental plasticity, context dependence, and organism-level embodiment of caste expression, while integrating these phenomena into a coherent realist ontology. Rather than presupposing that ‘castes’ correspond to autonomous biological units or fixed natural kinds, this approach treats them as condition-sensitive biological potentials whose manifestation depends on developmental, physiological, social, and ecological contexts.

#### 3.5.1 Realizable entities in BFO

Within BFO, realizable entities are a subtype of specifically dependent continuants, which are entities that inhere in a bearer, persist through time, and have the potential to be realized in appropriate circumstances (Arp et al. 2015). These include roles, functions, and dispositions, each distinguished by the nature of the conditions under which they may be realized. Importantly, realizable entities are temporally extended and context-dependent, which means that they may exist even when not currently realized, and their manifestation is contingent upon specific triggering conditions.

Dispositions capture inherent causal potentials grounded in the physical structure of an entity (e.g., a bee’s stinger disposition to pierce tissue). Functions denote realizable capacities essential to the kind of thing an entity is, often explained through evolutionary or etiological accounts (e.g., a nurse bee’s hypopharyngeal glands enabling brood-feeding). Roles, by contrast, are context-dependent capacities that are not essential to the bearer’s biological kind but arise through its participation in specific social, ecological, or institutional contexts (e.g., an ant’s role as a forager, as a nurse, or as a replete).

For biological systems, the realizable category is especially important because many organismal capacities (e.g., fertility, behavior, nest maintenance, aggression, brood care) are not continuously manifested. Instead, they exist as latent potentials whose realization depends on life stage, hormonal state, colony demography, social interactions, and environmental conditions. Structural features may enable these realizations, but structure alone is insufficient to determine when, how, or whether realization occurs.

Realizable entities thus describe phenomena that are not independent material units, but condition-sensitive potentials rooted in the properties of organisms and activated through specific processes. This makes them especially suited for modeling biological phenomena that are developmentally contingent, socially regulated, and temporally variable, which is precisely the conditions under which ‘caste’ differentiation in social insects occurs. This framework, thus, provides an ontological foundation for modeling ‘castes’ of social insects in bio-ontologies, whose identity is rarely exhausted by static morphology and instead depends on latent capacities and context-driven realizations.

#### 3.5.2 ‘Castes’ as realizable entities

When viewed through the lens of BFO, ‘castes’ of social insects align most closely with the category of roles, although their realization may involve the interaction of roles with organism-level dispositions and functions. ‘Caste’ identity emerges from individual developmental trajectories that are regulated by colony-level processes, including nutritional provisioning, pheromonal signaling, epigenetic regulation, and social feed-back mechanisms. The resulting phenotypes (such as queens, workers, or soldiers) do not constitute autonomous biological units, but are realizations of context-dependent developmental and physiological potentials instantiated in organisms.

Under this interpretation, a “queen” is not a natural unit in its own right but an organism that bears a reproductive role, realized through physiological processes such as oviposition and ‘caste’-specific glandular activity. Likewise, a “worker” bears a constellation of possible roles, such as foraging, brood care, nest construction, or defense, each realized only when the organism engages in those specific activities. These roles may be sequential, overlapping, or suppressed depending on the ‘colony’s’ internal state and ecological conditions. Although individuals sharing a ‘caste’ label may resemble one another phenotypically, they do not together form a causally unified system independent of their individual bodies.

Modeling ‘castes’ as realizable entities naturally accounts for several empirically well-established features of social insect biology. First, it accommodates developmental plasticity: larvae may bear multiple unrealized developmental potentials, with ‘caste’ outcomes determined by social and environmental cues acting during post-embryonic development. Second, it captures contextual sensitivity: the same genotype may realize different ‘caste’-related roles depending on ‘colony’ state (e.g., queen presence or absence, demographic imbalance, or resource availability). Third, it explains cross-’colony’ and cross-taxon variability: the same ‘caste’ label (e.g., “worker” or “soldier”) may correspond to different morphological, physiological, or behavioral profiles across species, because what unifies them is not an essential structure but a role realized within a particular social system. Fourth, it incorporates temporal dynamics: individuals may realize different roles over their lifespan, as seen in age polyethism in *Apis mellifera*, task switching in ants, or dominance-based reproductive transitions in some ponerine ants.

Importantly, this realizable entity model accommodates both morphologically canalized and morphologically plastic ‘caste’ systems. In species with developmentally fixed ‘castes’, structural traits constrain which roles can be realized, but realization remains temporally and contextually contingent. In more plastic systems, the same morphology may bear multiple unrealized role potentials, with ‘caste’ expression emerging from social regulation rather than predetermined form. In both cases, ‘caste’ identity is grounded in the realization of biological capacities rather than in membership in a natural kind.

These features are straightforward to represent using realizable entities but are incompatible with treating ‘castes’ as natural kinds or natural units. Modeling ‘castes’ as roles avoids the difficulties associated with intensional genus–differentia definitions, since roles do not require universal essential properties. Instead, role identity is determined relationally, meaning that they derive by the contexts in which the bearer participates and the processes it can realize.

By modeling ‘castes’ as realizable entities, specifically as roles instantiated in organisms under context-dependent developmental and social regimes, we obtain a flexible and ontologically coherent representation that aligns with both biological knowledge and BFO’s realist commitments. It provides a principled basis for representing ‘caste’-related phenomena in bio-ontologies and prepares the ground for more precise modeling of ‘caste’-associated functions, dispositions, and processes in social insect systems.

## 4 Discussion

### 4.1 Implications for biological representation and ontology design

Bio-ontologies play a central role not only in the standardization of data and meta-data, but also in data integration, interoperability, comparability, and long-term data management (Vogt et al. 2011). However, many existing ontologies have been idealized with highly specific practical goals in mind (Silva & Feitosa 2019a), often prioritizing the representation of narrowly defined domain entities. As a consequence, general ontological commitments concerning core biological categories are frequently left implicit or underspecified. This situation can lead to ontological inconsistencies, cross-ontology incompatibilities, and reduced semantic interoperability, ultimately undermining FAIR data principles. Adopting a shared upper-level framework, such as BFO, provides a principled foundation for aligning domain ontologies and improving interoperability across biological knowledge bases. It is important to note that the appeal to multiple frames of reference in this context is epistemic rather than metaphysical: different frames track distinct, real causal patterns in biological systems, but they do not themselves constitute additional kinds of entities. Ontological realism is preserved by grounding categories in mind-independent causal structures, even when those structures are accessed through heterogeneous representational perspectives.

Within this framework, the present analysis motivates a reassessment of how key entities in social insect biology (specifically ‘colonies’ and ‘castes’) are represented. Importantly, the role-based interpretation of ‘castes’ advanced here is not presented as the only conceivable ontological strategy for representing social differentiation in insects. Rather, it is defended as the only strategy that simultaneously satisfies three constraints imposed by realist ontology design under BFO: (i) avoiding the reification of context-dependent biological phenomena as natural kinds, (ii) accommodating temporal and developmental variability without category instability, and (iii) remaining fully compatible with existing representations of functions, dispositions, and processes. Alternative approaches — such as modeling ‘castes’ as processual classes, historically indexed phenotypic groupings, or functionally defined object aggregates — either collapse distinct frames of reference or reintroduce the same classificatory tensions that motivate the present reinterpretation.

By modeling ‘colonies’ as unified causal systems and natural units, and ‘castes’ as realizable entities instantiated in individual organisms, the proposed approach enforces clear ontological distinctions between independent biological individuals and context-dependent biological capacities. Importantly, this modeling strategy remains robust in the face of ongoing empirical discoveries. As new insights into ‘caste’ determination, developmental regulation, or evolutionary transitions emerge, particularly from eco-evo-devo research and high-throughput genomic and epigenomic studies, the realizable-entity framework allows concepts to be refined, extended, or reinterpreted without requiring wholesale revision of the underlying ontology. This flexibility is especially valuable given the substantial knowledge gaps that remain regarding the proximate and ultimate drivers of ‘caste’ differentiation.

Although biological representations are necessarily mediated by cognitive and linguistic practices, ontological realist commitments aim to represent aspects of biological reality as mind-independent entities. In the present framework, this is achieved by demarcating entities according to multiple frames of reference beyond the spatiostructural one. ‘Colonies’ are identified through causal unity and historical continuity, whereas ‘castes’ are captured through developmental, functional, and social frames of reference. Explicitly grounding these distinctions in relations of causal unity helps prevent category mistakes, such as reifying context-dependent developmental outcomes as autonomous biological individuals. Moreover, this strategy encourages ontology developers, with the intimate support of biologists, to first identify the causal and regulatory properties that confer connectedness and stability on biological entities, and only then to formulate operational definitions and classification criteria.

Formalizing ‘colonies’ and ‘castes’ in this manner also has implications for long-standing debates concerning downward causation in biological systems (Craver & Bechtel 2007). While all members of a colony typically share a common genetic background, ‘caste’ differentiation arises through differential gene expression mediated by ‘colony’-level regulatory processes (Crozier & Pamilo 1996; Elsner et al. 2018). Representing ‘colonies’ as higher-level causal systems and ‘castes’ as realizable entities borne by organisms provides a clear ontological scaffold for modeling how ‘colony’-level states constrain, enable, or modulate organism-level developmental trajectories without invoking metaphysically problematic forms of causation.

From an ontology-design perspective, recent efforts to enrich relation vocabularies [such as the introduction of developmentally-related-to relations (Buttigieg et al. 2016; Walls & Buttigieg 2017)] offer useful tools for representing complex biological dependencies. However, attempting to organize ‘caste’ categories solely through class–subclass hierarchies risks fragmenting ontologies into multiple, frame-specific taxonomies. As noted by Vogt et al. (2012), this problem can be mitigated by distinguishing between different kinds of taxonomic relations (e.g., physiological, developmental, or functional) and by relying on logical definitions and automated reasoning rather than rigid single-inheritance trees.

Within BFO, modeling ‘castes’ as realizable entities (specifically as roles) provides a coherent alternative. Roles are inherently context-dependent and process-realized, making them well suited for representing temporal polyethism, task flexibility, and environmentally mediated developmental outcomes. As realizable entities, such roles are temporally qualified, i.e., an organism may bear a given ‘caste’-related role at a particular time and cease to bear it at another, depending on developmental state, colony-level conditions, or environmental context, without undergoing any change in its identity as an organism. This approach dissolves the tension between structural and dispositional accounts of ‘caste’ differentiation: morphological traits constrain which roles can be realized, but they do not exhaust ‘caste’ identity. Consequently, both morphologically fixed systems and plastic systems can be represented within a single, unified ontological framework.

Notably, this role-based interpretation is compatible with diverse forms of social organization, including systems with irreversible morphological differentiation (e.g., many ants), systems with early, morphologically canalized ‘caste’ differentiation (e.g., termites), and systems in which queens and workers are morphologically distinct but worker task allocation remains developmentally and temporally flexible (e.g., social bees), as well as systems exhibiting extensive temporal polyethism across individuals. In all cases, ‘caste’ identity is modeled independently of any single developmental or morphological criterion.

From a formal perspective, this interpretation suggests the following modeling strategy. ‘Caste’ would be represented as a subclass of *role* (BFO:0000023), inhering in individual organisms and realized in biological processes. For example, the basic subclass structure can be expressed as follows.

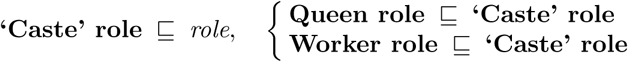

With axioms such as:

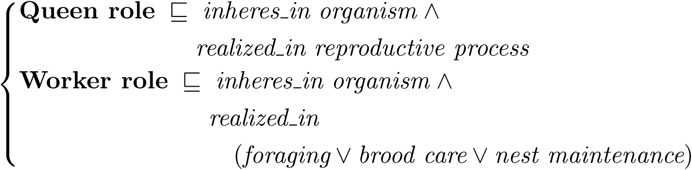

Under this scheme, organisms may bear multiple ‘caste’-related roles over time, and the realization of these roles can be conditioned on colony state, developmental stage, or environmental context, without requiring reclassification of the organism itself.

### 4.2 Current limitations for ‘caste’ subcategories

In groups that express intermediate or phenotypic mosaics or have dispositions to perform specific functions or roles within a ‘colonial’ environment (such as foraging or brood-care), it is common to establish a subcategory of ‘caste’, i.e., ‘subcaste’ (Silva & Feitosa 2019a). In pragmatic cases – i.e., when researchers have specific investigative motivations for splitting ‘caste’ categories – it would be possible to define some operational criteria, based on the respective frame of reference, to further classify ‘castes’ into ‘subcastes’. However, if we wished to argue that ‘subcastes’ truly represent a subclass of ‘castes’, the former would necessarily need to inherit all properties of the latter and incorporate additional defining properties, thus narrowing the meaning of the term. However, insofar, attempts to split ‘castes’ into subcategories have not been entirely successful. This difficulty is both empirical and ontological: if ‘castes’ themselves lack stable defining properties and causal unity, then attempts to subdivide them into ‘subcastes’ inherit and amplify these limitations.

One prominent example refers to the further categorization of ‘castes’ proposed by Wilson (1979). According to Wilson’s proposition, a ‘caste’ would be an ensemble of colony members that specialize in particular tasks for prolonged periods, being typically (but not necessarily) distinguished by other genetic, anatomical, or physiological traits. He further categorizes ‘castes’ into two subcategories, namely ‘physical castes’ and ‘temporal castes’, with the former being defined by allometric and other anatomic criteria and the latter being defined by age grouping. It is noteworthy to highlight that, although being operationally sound for the most part, the translation of Wilson’s proposition to bio-ontologies has some limitations: (i) ambiguous defining criteria of ‘castes’, which are subsequently used to derive the defining criteria of the proposed subcategories; and (ii) epistemic and representational difficulties in specifying recognition criteria for functionally defined caste categories. A further consequence of these issues concerns the apparent unnestedness of caste-related categories when translated into formal classificatory hierarchies.

Wilson’s first issue directly stems from the lack of consideration of the distinct frames of reference used to define the recognition criteria for the class being demarcated. Wilson considers ‘castes’ as object aggregates in the sense of an organism as an aggregate of cells and molecules that are recognized by a set of functionally defined spatiostructural entities (i.e., anatomic distinct individuals that have the disposition to perform specific biological functions). However, recognition criteria of functionally defined spatiostructural entities cannot be directly inferred from their defining properties (i.e., dispositions), because said properties are not usually unambiguously bound to a specific set of spatiostructural properties (Vogt et al. 2022). According to Peeters (2012), certain groups of social insects possess this characteristic uncoupling of disposition (in his example, reproductive disposition) from a specific set of spatiostructural properties (e.g., development of morphoanatomical components associated with wings). In cases where researchers want to find specifiable spatiostructural recognition criteria for functionally defined biological entities, they must conduct experiments that can identify a particular functionally defined spatiostructural entity (e.g., through physiological experiments). Alternatively, researchers could create a context in which a reaction is triggered that realizes the disposition and the reaction has directly or indirectly observable consequences, which can be used as recognition criteria. This leads to Wilson’s ”second issue”.

Although these issues appear merely as terminological, they also have direct consequences for ontology design, particularly when attempting to formalize ‘subcastes’ within hierarchical classification systems.

One very important thing to highlight is that most of the time, researchers do not have access to these experiments, and, when they try to specify recognition criteria for functionally defined spatiostructural entities, they are restricted to differential diagnosis and comparative approaches coupled with conclusions based on analogy (Vogt et al. 2022). As a compound, non-deductive, type of reasoning (Minnameier 2010; Peirce 1896: 28p, CP 1.65), analogical reasoning involves inferential moves that spam different frames of reference – i.e., it involves processes of matching and mapping between different conceptual domains – which can lead to logical inconsistencies when misapplied. However, despite the complexity of the inferential form of analogy, one can recognize ‘castes’ by a set of functionally defined spatiostructural entities without potentially leading to ontological errors. Additionally, in terms of ontology design, unnestedness does not necessarily lead to errors; rather, the strict adherence to single-inheritance hierarchies when dealing with compositional classes can potentially lead to errors. In this sense, what might appear as a third issue — namely, the apparent unnestedness of caste-related categories when translated into classificatory hierarchies — is better understood as a design consequence of the preceding epistemic and ontological issues, rather than as an independent problem in ontological requirements. As a solution for the single-inheritance hierarchy question, researchers can account for additional class-subclass relationships by specifying within class axioms some properties that its instances necessarily have to possess and then use these axioms to infer subclasses. This way, it is possible to have a single-inheritance ontology - in which each class has only one direct parent class - and derive all implicit class-subclass relationships through reasoning.

Taken together, these considerations suggest that difficulties in defining ‘subcastes’ arise not from insufficient empirical resolution, but from treating context-dependent realizable phenomena as if they were stable taxonomic kinds, which is an issue that can only be resolved by rethinking how caste-related distinctions are represented in ontological models.

## 5 Conclusion

Using a well-defined ontological framework to represent ‘colonies’ of social insects as natural units enables a coherent and realist representation of multiple dimensions of biological organization. By treating ‘colonies’ as entities unified through relations of causal connectedness and historical continuity, this approach provides clear criteria for partitioning social insect systems into identifiable individuals. Although future work is needed to resolve specific modeling choices in bio-ontologies, such as whether ‘colonies’ should be represented as instances of a fine-grained class (e.g., *’Colony’ of social insect* ) or as instances of a more general class like BFO’s *object aggregate*, the adoption of a logically aligned framework constitutes a substantial step toward resolving persistent inconsistencies in the demarcation of fundamental entities in social insect biology.

Reinterpreting ‘castes’ as realizable entities rather than as natural kinds provides a more ontologically consistent and biologically faithful account of social organization in insects. By modeling ‘caste’ as a role borne by individual organisms and realized in specific biological processes, this framework accommodates temporal variation, developmental plasticity, and context dependence without forcing reclassification of organisms themselves. This shift resolves long-standing conceptual ambiguities in ‘caste’ representation, aligns ontological commitments with contemporary biological understanding, and offers a robust foundation for the logical formalization of social categories in bio-ontologies.

More broadly, the distinctions articulated here illustrate how careful separation between natural units and realizable entities can improve the representation of complex social systems in biology, with implications extending beyond social insects to other domains where context-dependent biological organization plays a central explanatory role.

## Acknowledgments

I am deeply grateful to Lars Vogt, who pointed out several inconsistencies and misconceptions in previous versions of this work and proposed solutions to conflicts raised in the discussion.

## Authors’ contributions

Conceptualization, methodology, visualization, writing, reviewing and editing were all made by TSRS.

## Data Availability

All data is available in the manuscript.

## Ethical approval

This article does not contain any studies with human participants or animals, performed by any of the authors.

## Consent to participate

All authors agree to participate in this research project.

## Consent for Publication

All authors approve the publication of the manuscript.

